# Insights on protein thermal stability: a graph representation of molecular interactions

**DOI:** 10.1101/354266

**Authors:** Mattia Miotto, Pier Paolo Olimpieri, Lorenzo Di Rienzo, Francesco Ambrosetti, Pietro Corsi, Rosalba Lepore, Gian Gaetano Tartaglia, Edoardo Milanetti

## Abstract

Understanding the molecular mechanisms of thermal stability is a challenge in protein biology. Indeed, knowing the temperature at which proteins are stable has important theoretical implications, which are intimately linked with properties of the native fold, and a wide range of potential applications from drug design to the optimization of enzyme activity.

Here, we present a novel graph-theoretical framework to assess thermal stability based on the structure without any a *priori* information. In our approach we describe proteins as energy-weighted graphs and compare them using ensembles of interaction networks. Investigating the position of specific interactions within the 3D native structure, we developed a parameter-free network descriptor that permits to distinguish thermostable and mesostable proteins with an accuracy of 76% and Area Under the Roc Curve of 78%.

## Introduction

Temperature is one of most crucial factors organisms have to deal with in adapting to extreme environments^1^ and plays a key role in many complex physiological mechanisms^2^. Indeed a fundamental requirement to ensure life at high temperatures is that the organisms maintain functional and correctly folded proteins^2–4^. Accordingly, evolution shapes energetic and structural placement of each residue-residue interaction for the whole protein to withstand thermal stress.

Studying thermostability is fundamental for several reasons ranging from theoretical to applicative aspects^5^, such as gaining insight on the physical and chemical principles governing protein folding^6–8^, and improving the thermal resistance of enzymes to speed up chemical reactions in biopharmaceutical and biotechnological processes^9,10^.

Despite the strong interest in thermostability^11–13^, its prediction remains an open problem. A common descriptor used to quantify the thermal stability of proteins is the melting temperature (T_m_), defined as the temperature at which the concentration of the protein in its folded state equals the concentration in the unfolded state. To date, computational approaches, both sequence- and structure-based, have exploited statistical analysis^8,14,15^, molecular dynamics^16,17^ and machine learning^18,19^ to predict the melting temperature. Most of the studies are based on comparative analyses between pairs of homologues belonging to organisms of different thermophilicity^20,21^.

Predicting the stability of a protein *ab initio* using a structure based-approach has never been achieved so far. Lack of success in this area is mostly due to limitations in our knowledge about the relationship between thermal resistance and role of the interactions that stabilize a protein structure^22^. Some differences in terms of amino acid composition or spatial arrangement of residues have been reported^8,23–25^. One of most notable differences involves the salt bridges: hyperthermostable proteins have stronger electrostatic interactions than their mesostable counterparts^26^. Recently Folch *et al*.^22,27^ reported that distinct salt bridges may be differently affected by the temperature and this might influence the geometry of these interactions as well as the compactness of the protein. Core packing seems related to thermal resistance at least to some extent^28^. Yet, a lower number of cavities and a higher average relative contact order (i.e. a measure of non-adjacent amino acid proximity within a folded protein) have been also observed while comparing thermostable proteins with their mesostable paralogs and orthologs^6^. Noteworthy, the hydrophobic effect and residue hydrophobicity seem to play a rather marginal role on protein stabilization^29–31^, while they are considered the main forces driving protein folding.

Here, we present a new analysis based on the graph theory that allows us to reveal important characteristics of the energetic reorganization of intramolecular contacts between mesostable and thermostable proteins. In light of our results and to promote their application, we have designed a new computational method able to classify each protein as thermostable or as mesostable without using other information except for the three-dimensional structure.

## Results

### Uncovering the differences in energetic organization

Aiming at the comprehension of the basic mechanisms that allow proteins to remain functional at high temperature, we focused on the non-bonded interactions between residues that play a stabilizing role in structural organization^32^. In particular, we investigated how different thermal properties are influenced by the energy distribution at different layers of structural organization. We analyzed the interactions occurring in proteins of the *T_whole_* dataset, containing the union of the T_m_ dataset (proteins with known melting temperature taken from ProTherm database^33^) and the *T_hyper_* dataset (proteins belonging to hyperthermophilic organisms, with *T_env_* ≥ 90*^o^C* collected from Protein Data Bank^34^), for a total number of 84 proteins (see Methods). To describe the role of single residues in the complex connectivity of whole protein, we adopted a graph theory approach describing each protein by the Residue Interaction Network (RIN): each residue is represented as a node and links between residues are weighed with non-bonded energies (as described in Methods).

At first, we investigated the relationship between thermostability and energy distribution of intramolecular interactions. To this end, the *T_m_* dataset was divided into eight groups according to protein *T_m_* and for each group the energy distribution was evaluated, as shown in Figure 1a. The general shape of the density functions is almost identical between the eight cases, independently from the thermal properties of the macromolecules, and this is clearly due to the general folding energetic requirements.

**Figure 1.**
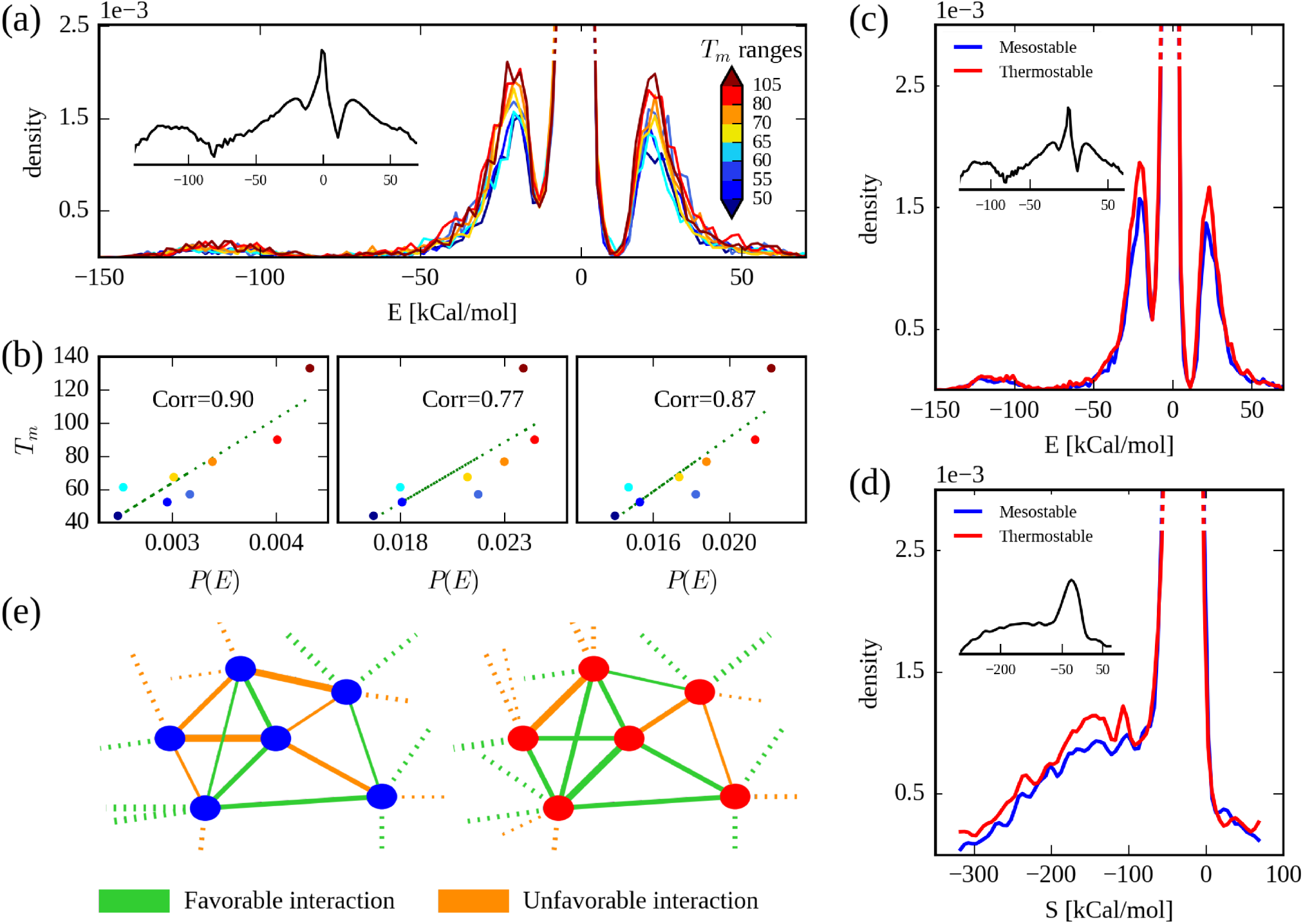
(color online) **a)** Probability density distributions of total interaction energies for the eight subsets defined in the *T_m_* dataset. Each distribution is built using a group of proteins whose melting temperatures lie in the same range. The eight ranges, from lower to higher *T_m_*, are represented by colors from dark blue to dark red. The density functions exhibit a dependence with the melting temperatures ranges and peak heights increase with the temperatures. Inset shows the energy distribution in log-scale obtained using all proteins. **b)** Correlation between the area of each density peak and the average *T_m_* for the eight dataset groups. **c)** Probability density distributions in log-scale of total interaction energies for mesostable (blue) and thermostable (red) proteins belonging to the *T_whole_* dataset. Inset shows the energy distribution in log-scale obtained using all proteins. **d)** Probability density distributions in log-scale of Strength network parameter for mesostable (blue) and thermostable (red) proteins belonging to the *T_whole_* dataset. Inset shows the Strength distribution in log-scale obtained using all proteins. **e)** Schematic representation of the strong favorable and unfavorable interactions both for a mesostable (left) and a thermostable network (right).

A strong dependence between thermal stability and the percentage of strong interactions is evident looking at the disposition of the density curves (Figure 1a): the higher the thermal stability the higher the probability of finding strong interactions. Yet, less thermostable proteins possess a larger number of weak interactions. In particular, as shown in Figures 1a-c, it is possible to identify three ranges of energies that correspond to three peaks of probability density, i.e. a very strong favorable energy region (*E* < −70 kCal/mol), a strong favorable energy region between −70 kCal/mol and −13 kCal/mol, and a strong unfavorable interaction region (*E* > 11 kCal/mol). More formally, for a protein the probability of having an interaction with energy E, *P*(*E*), in the three ranges linearly depends on the protein melting temperature with correlation coefficients of 0.90, 0.85, 0.87, respectively (Figure 1b).

In order to have strong-signal sets, we reduced the division in just two groups, classifying proteins as mesostable or thermostable if their melting temperatures are, respectively, lower or higher than 70*^o^C*, which is the optimal reaction temperature of thermophilic enzymes^35–40^. In this way the energy distributions in Figure 1a are calculated only for the mesostable and thermostable distributions in Figure 1c (*T_whole_* dataset). The two-group division allows us to include the hyperthermophilic proteins in our analysis, since their *T_m_* is higher than the threshold. The two resulting distributions, found to be significantly different with a p-value of 4.2 × 10^−46^ (nonparametric test of Kolmogorov-Smirnov^41^), have an expected value at −0.5 kCal/mol and negative interactions have a probability of more than 60% to be found. Regions below −13 kCal/mol and above 11 kCal/mol represent the 6.6% and 5.2% of the total energy for thermostable and mesostable proteins respectively. Typically, such energies require the presence of at least one polar or charged amino acid and in particular Arg, Asp, Glu and Lys are involved in more than 90% of the interactions. Noteworthy, the small fraction of energies centered near −120 kCal/mol (see Figure 1a) are due to polar or charged amino acid interactions taking place at short distance. Next, we investigated the residue Strength, defined for each node as the sum of the weights of its links (see Methods). The two Strength distributions for mesostable and thermostable proteins are shown in Figure 1d. Even in this case, they are different according to Kolmogorov-Smirnov test with a p-value of 1.9 × 10^−9^.

For the first time, our analysis provides both a general intuition on the protein folding and a specific insight on thermal stability. Even if strong positive and strong negative peaks have a comparable height (Figure 1c), the rearrangement of protein side chains masks the positive interactions, substantially preventing the condensation of unfavorable interactions in a single residue, as testified by the small probability of finding a residue with a positive Strength. Indeed, for the whole dataset, there is more than 97% of probability of finding a residue with negative Strength. The most frequent value is found at −27 kCal/mol, with a change in the slope of the density functions around −70 kCal/mol and 5 kCal/mol, corresponding to the regions with negative and positive Strengths. At the Strength level of organization, a difference between thermostable and mesostable proteins is found. Indeed, residues belonging to the group of thermostable proteins show a higher probability of having high negative Strength values with respect to the mesostable ones, testifying an overall higher compactness of thermostable protein fold. In particular, the probability of finding a node having a Strength below −70 kCal/mol is 19.9% and 16.1% for thermostable and mesostable respectively.

Figure 1e shows a schematic representation of the organization of strong energies both for mesostable proteins and thermostable proteins. In fact, the most important finding of our analyses is that thermostable proteins have more favorable energies concentrated in a few specific residues. In contrast, mesostable proteins tend to have a less organized negative residue-residue interactions network. Given this different way to rearrange amino acidic side chains between proteins with different thermal properties, we mapped the energetic distribution on the protein secondary structures in order to study how energetic allocation is reflected on a higher level of organization.

To do so, we retrieved the secondary structures (helices, strands, loops) for all the proteins of the *T_m_* dataset using DSSP^42^ and assigned each residue-residue interaction occurring in a protein to six possible classes (helix-helix, helix-strand, helix-loop, strand-strand, strand-loop and loop-loop), according to the secondary structure residues belong to (see Methods).

The goal of the analysis is to determine whether there is a secondary structure class containing more energy than one would expect by chance. To this end, we estimated the difference in energy of a specific class with respect to the average energy (i.e. energy that the same class would had if uniformly assigned). In the group of mesostable proteins, pairing between residues of the same structure (helix-helix, strand-strand and loop-loop) is associated with higher-than-average energies, while mixed combinations (helix-strand, loop-strand and loop-helix) have lower energies. As shown in Figure 2a, a surplus is committed to helix-helix interactions, while helix-strand has a negative difference of about 8.5%. When the thermal resistance is taken into account the most significant distinction between the mesostable and thermostable groups is in the helix-loop pairing. Thermostable proteins have a larger shift than the mesostable counterparts. These results suggest a stabilizing role of helix-loop interactions and we argue that thermostable proteins preferentially gather their energy to this specific class (Figure 2b).

**Figure 2.**
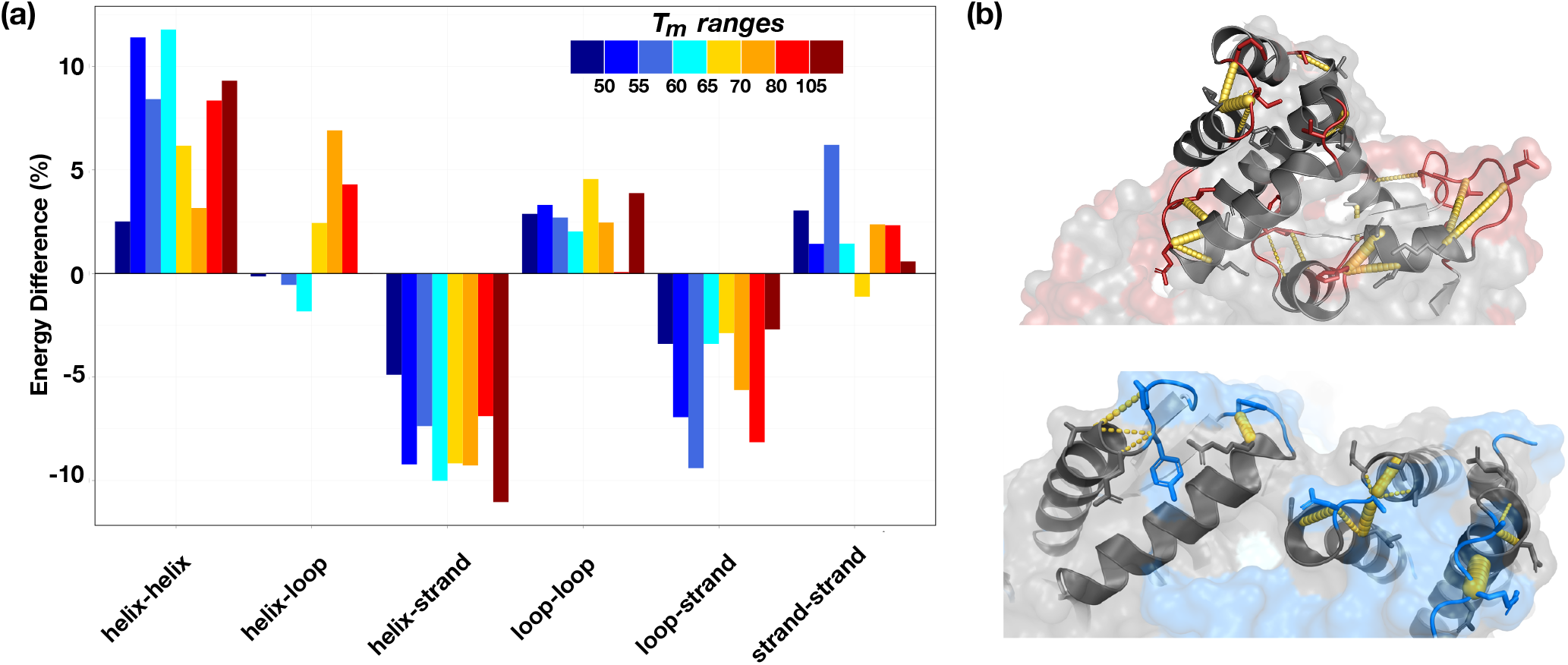
**a)** (color online) For each class of interaction, we report the difference in percentage between actual energy and the expected value of the specific group assuming an uniform distribution of energies. **b)** Cartoon representation showing a region of a thermostable (top) and mesostable (bottom) protein. Strong interactions between charged-charged amino acid belonging to loop-helix secondary structure are shown in yellow sticks. Loop secondary structures are indicated in red for the thermostable protein and in blue in the mesostable one. All the interactions are represented in yellow.

### Assessing protein thermal stability

In the light of our findings on the difference on the energy organization between mesostable and thermostable proteins, we looked for a way to assess the thermal resistance of a protein given its structure. The simplest way to quantify the impact of energy distribution on the thermal resistance is the comparison with a protein of same structure but different energy organization, i.e. a homologue (and indeed this has been widely done^43^). Ideally, differences between two homologous proteins with different thermal stability are attributable only to their different thermal resistance. The more pronounced reorganization of the interactions in thermostable proteins confirms that they undergo an evolutionary optimization process which introduces fold-independent correlations in the spatial distribution of the interactions. By contrast, mesostable proteins have not these correlations, thus with respect to thermal stability, their energy organization can be considered more random.

We designed a procedure that compares a given protein with modified versions of itself where protein structure is preserved, while chemical interactions have energies typical of mesostable proteins and randomly assigned in a physical way, i.e. maintaining residue-residue distance information (see Methods). In this way, the randomization strategy provides a way to compare each real protein network with an ensemble of re-weighted cases, having the same number of nodes and links but with new weights (i.e. energies). These energies are extracted from the mesostable energy distribution using the interaction distance as constraint for the sampling. This procedure has the purpose of disrupting the evolutionary optimization and it is expected to have a larger effect on the highly organized network of thermostable proteins. By virtue of the different energy distribution between mesostable and thermostable proteins, sampling mesostable energies allows to properly assess the difference between the real thermostable protein network and its randomized counterpart. All steps of the randomization process are schematically illustrated in Figure 3. In particular, given a link characterized by an energy weight *E_ij_* and by a distance of interaction *d_ij_*, we replaced the energy with a new one 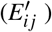 extracted from a energy distribution defined for the specific distance interval *d_ij_* belongs to. For each distance interval k, we generated a probability density function *ρ_k_*(*E*), using only the energies values observed in such interval in the mesostable proteins. At the end of the process, for each real RIN, we generated an ensamble of random networks (rRINs). The randomization allows us to develop a classifier based on the distance between the real network Strength and the random Strength distribution. The *T_s_* score, defined in Eq. 7 (see Methods), is a measure of how much the original RIN average Strength value deviates from the expected average value of the rRIN distribution. Note that our descriptor is general and parameter-free and can be computed for every kind of weighted graph. The *T_s_* score can be used as a thermal stability classifier setting the threshold value at 0; substantially considering true all predictions for which the *T_s_* score is higher than 0 and the protein *T_m_* is higher than 70*^o^C* or alternatively the *T_s_*score is lower than 0 and the protein *T_m_* is lower than 70*^o^C*. A so defined method is completely parameter-free. It only requires a probability density of mesostable protein interactions. In order to evaluate a possible dependence of the method from the chosen dataset, we performed a cross-validation (7-folds see Method) using the *T_s_* score computed with total energy Strength. The method achieves an average accuracy of 72% plus or minus 3% with a mean ROC curve characterized by an AUC value of 80% plus or minus 2%. The small error on both the performances (due to the dimensions of the dataset) indicates the independence of the method from the input information.

**Figure 3.**
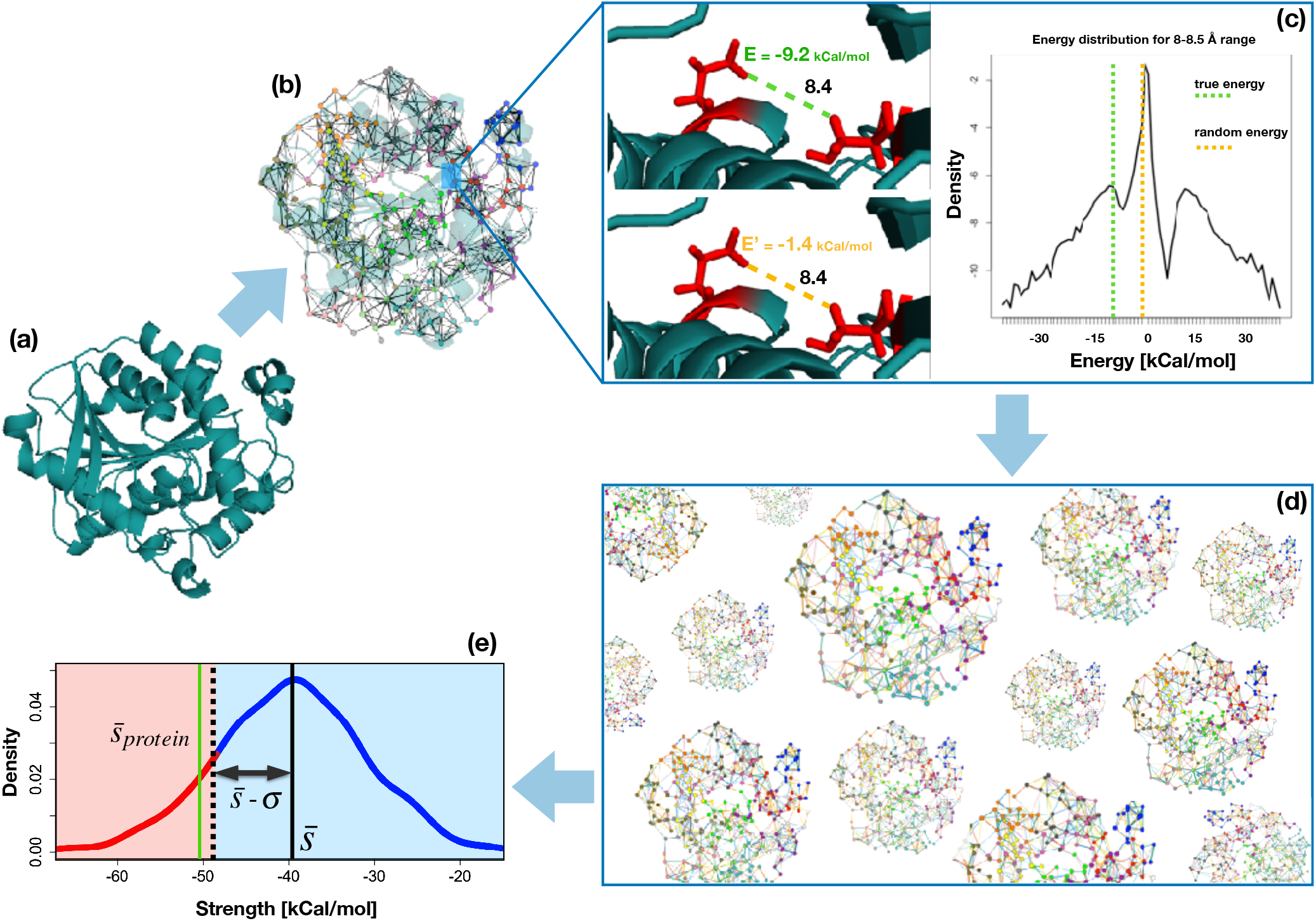
(color online) Given a protein structure **(a)**, our method represents it as a Residue Interaction Network **(b)**, where each amino acid becomes a node and the energetic interactions between amino acids are weighted links connecting the nodes, **(c)** The first step of the procedure is to calculate the minimal atom-atom distance for each residue pair, which is indicated in yellow on the left of panel. In the example here, the minimal distance between the two residues is 8.4 Angstrom. The energy value related *to* such contact is replaced with another one, randomly extracted from the energy distribution of mesostable proteim derived only considering energies in the distance interval which corresponds to the minimal distance of the specific eontact. In the middle of panel the density of energy belonging to the dietance range 8-8.5 Angstrom is shown. The news energy is represented with the green line in the right of ihe panel. Performing this procedure for each pair, a new network of intramolecular interactions is established characterized by new energy organization. Reiteraiing the process many times, we obtain an ensamble of random networks **(d)**. Finally, for each random network the average Strength parameter is calculated. Panel **(e)** shows the Strength distribution obtained iterating the procedure. Green line represents the mean Strength value of the real network, while red and blue region in the random Strength distribution show the classification criterion: if real Strength lies in red region the protein is classified as thermostable while if it sets in the blue region it will be labeled as mesostable.

Classifying on the basis of the 0 threshold of the *T_s_* score, i.e. considering the *T_s_* as a binary variable, loses part of the information contained in the descriptor. In order to have a more sensible classification, we evaluated three different scores, using the total energy (defined in Eq. 5) and specific interaction terms, i.e. the Coulomb and Lennard-Jones interactions (Eq. 3 and Eq. 4), and performed a clustering analysis. Figure 4(a) shows the hierarchical clustering obtained clustering all the proteins of our *T_whole_* dataset using the Ward method as linkage function while the Manhattan distance among the three descriptors was used as distance metric. We also tested different metrics and clustering methods obtaining very similar results (data not shown). The optimal clustering cut was estimated using both the Connectivity, Dunn and Silhouette parameters, which indicates the two group division as the optimal one. We called these groups “Mesostable” (right group in Figure 4) and “Thermostable” (left group). Indeed, the right cluster, containing 47 proteins, includes almost exclusively mesostable proteins (38), while the left cluster contains 26 thermostable proteins over the total 37 proteins. The overall accuracy of the method is 76%. We correctly assign the right thermal stability to 64 out of 84 proteins. The AUC of the ROC curve for the three *T_s_* descriptors are 0.78, 0.79 and 0.68 (Figure 5a).

**Figure 4.**
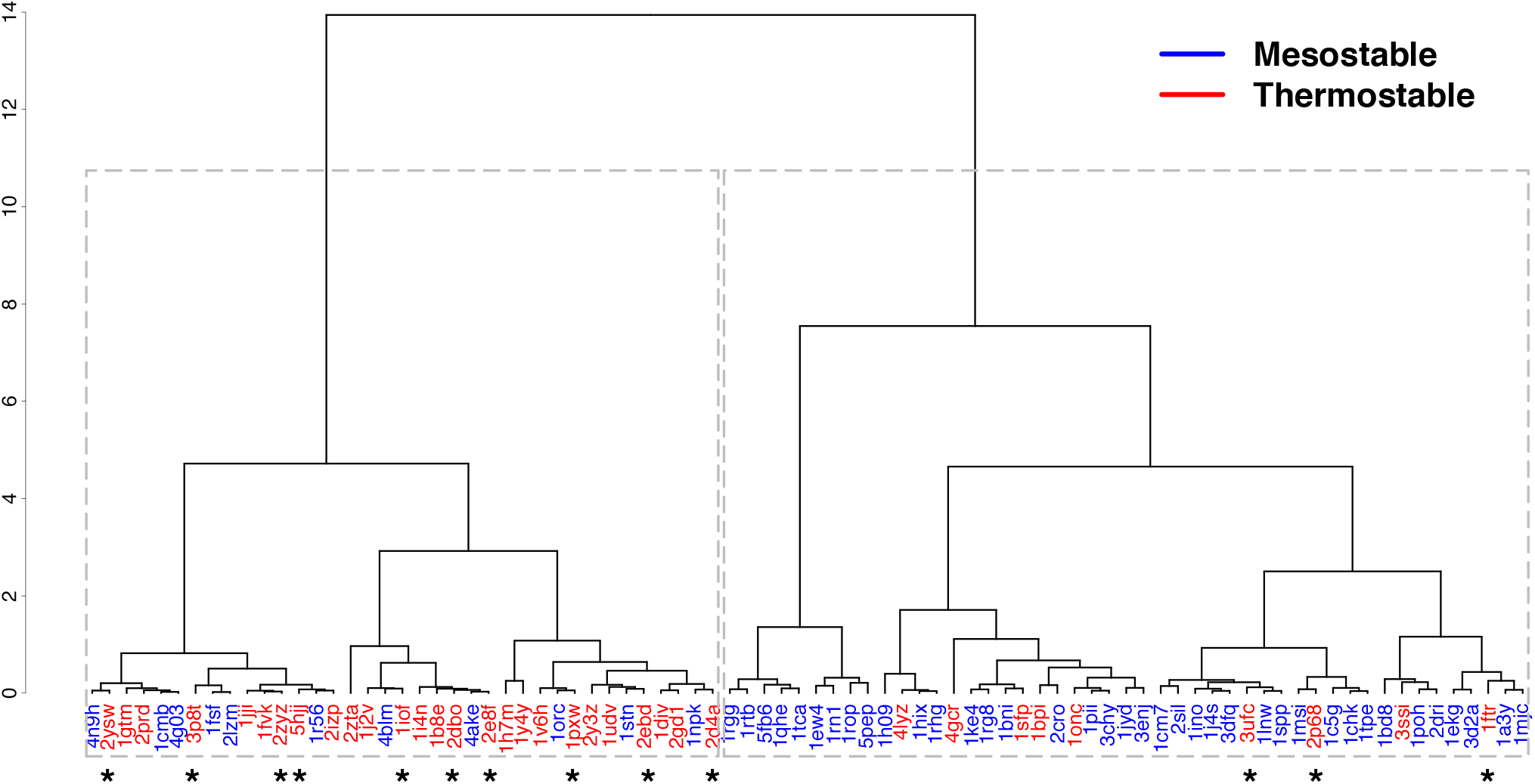
(color online) Cluster of the *T_whole_* dataset proteins with three Strength based descriptors, i.e. Coulomb, Lennard-Jones and total energy. Stars indicate proteins on the *T_hyper_* dataset. The two groups are discriminated with a p-value of 2.6 × 10^−6^ (Fisher’s exact test).

### Key residues identification

Here, we investigated the thermal resistance properties of proteins at the residue level. As protein stability is the result of the cooperative effects and the synergic actions of several residues, assessing the specific contribution of each amino acid is difficult^44^. We define the 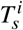 score for each residue according to Eq. 8, creating two groups of residues for each protein: with 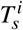 lower or higher than zero. We consider residues belonging to the first group to have a more stabilizing role than the ones in the second group. Consequently, along the lines of the global-protein classification procedure, we defined “thermostable” (respectively “mesostable”) residues belonging to the first (second) group. Using a total-energy based score, thermostable residues are the (11 ± 4) % of total residues.

As expected, thermostable proteins have more thermostable residues with respect to mesostable ones (12% compared to 9%). Furthermore, by repeating the same analysis using Coulomb (C) and van der Waals (vdW)-based scores, we found that the average number of thermostable residues is the 11% and 16% of total residues, respectively. In the Coulomb network (see Figure 6a), the most frequent thermostable amino acids are the four charged amino acids: Arg, Asp, Glu and Lys, which cover the 96.6% and 96.1% of thermostable residues in thermostable and mesostable proteins respectively.

Apolar and aromatic residues (Leu, Met, Phe and Tyr) are typically thermostable residues of the van der Waals network, including 53% and 54% of the total residues in mesostable and thermostable proteins, respectively (see Figure 6b).

In order to investigate the role of each residue in the complexity of the whole system, we analyzed the properties of all residues using a graph-theory approach, calculating 8 network parameters, i.e. Betweenness Centrality, Closeness Centrality, Strength, Diversity Index, Mean Shortest Path, Hub Score, Clustering Coefficient and Degree (see Methods). A Principal

Component Analysis (PCA) was performed in both kinds of network. In Figure 6c-d, all residues were projected along the first three principal components, which represent the 82% and 63% of the variance for van der Waals and Coulomb network respectively (see Supplementary Information). Thermostable residues are neatly separated from others if we consider the largest eigenvalue of the PCA in the Coulomb network and more weakly if we take into account the second and third ones. The Strength and the Closeness Centrality are the most relevant loadings along the first eigenvector both in favorable and unfavorable interactions. In vdW networks (Figure 6d), residue splitting in the PCA planes is less pronounced, even if a separation does occur along the second and third components, having the Clustering Coefficient and the Diversity Index as principal loadings (see Supplementary Information). In both kinds of network, independently from the protein of origin, thermostable residues populate the same regions of the PCA planes (green and orange dots in Figure6). Considering all dataset residues, about 25% of them are charged and only the 11% is classified as thermostable. A similar evidence was found for the four thermostable amino acids identified by vdW score. The role played by charged and aromatic amino acids in thermal resistance has been investigated in previous comparative studies^45,46^. Generally, charged residues form highly energetic electrostatic cages which prevent water inclusion^47,48^ while apolar and aromatic amino acids form short-ranged vdW interactions that confer stability to the overall structure^49,50^. Here we identify key residues whose peculiar spatial disposition confers them a particular role in the stabilization of the protein. Notably, our approach, based on an heterogeneous dataset, permits us to confirm and generalize the stabilizing role of both the charged and apolar/aromatic residues formerly suggested by homologous-based studies.

The mean shortest path (L) and the clustering coefficient (C) are able to catch the effect of the thermostable residues on maintaining these important structural motifs. The former provides information about the position of the residue in the network with the most central residues, having higher shortest path values. The latter quantifies the residue surrounding packing, being a ratio between the actual links and maximal number of possible links^51–53^.

In Figure 6e (left panel), we projected all residues in the LC plane coloring in dark red the charged thermostable residues and in cyan the charged non-thermostable residues. Charged residues are concentrated in the region characterized by both small L and C values, with their thermostable subset tending to possess the smaller possible value of C. This means that thermostable residues have both to be exposed and surrounded by residue that make low energetic interaction between each others. In analogy with coulombian networks, we projected in the LC plane the four kinds of key residues identified in the vdW networks. Even if the signal is weaker, key residues in the thermostable van der Waals network (Leu, Met, Phe e Tyr) tend to possess a higher clustering coefficient, testifying the packing stabilizing effect of vdW interactions. Densities of C parameter are found to be different with a p-value < 10^−16^ (nonparametric test of Kolmogorov-Smirnov).

These finding allow us to divide residues in 8 groups: four groups are identified by the Coulomb interaction, i.e. thermostable charged/uncharged residues and non-thermostable charged/uncharged residues; while vdW interaction networks divide residue according to thermostable/non-thermostable being or not being in the Leu-Met-Phe-Tyr group. For each protein of the *T_whole_* dataset it is possible to compute the sum of the 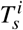 scores in each of the 8 possible groups, obtaining a vector of 8 descriptors for each protein. Performing a linear regression with the four Coulomb-based vector component, the four vdW-based ones and with the whole eight-component vector we end up with a preliminary AUC of the ROC curves of 0.81 e 0.77 and 0.83 respectively (see Figure 5b), and we are currently developing a residue-specific approach for *T_m_* prediction.

**Figure 5.**
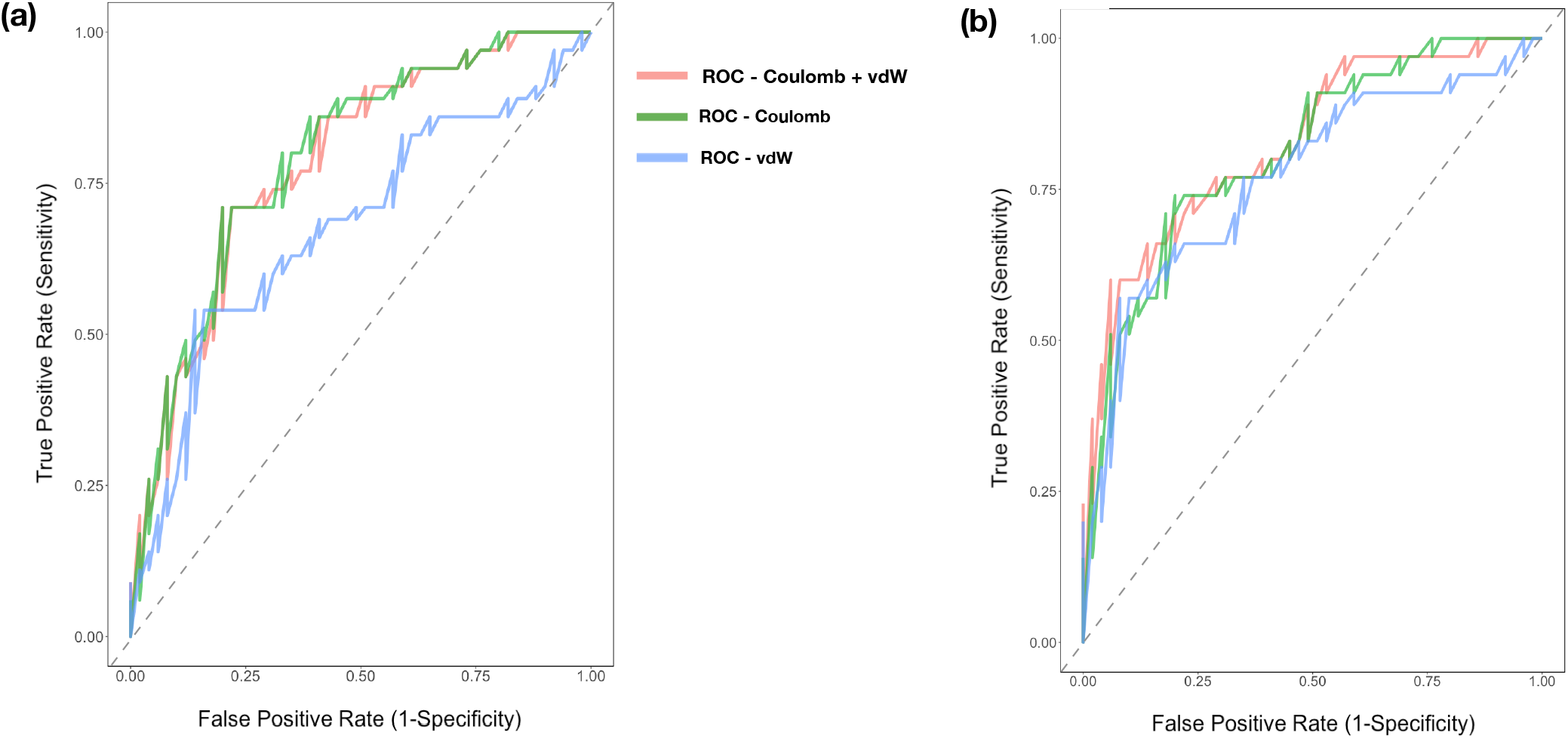
(color online) **a)** ROC curves of the three descriptors with the whole network *T_s_* scores. **b)** ROC curves of the three descriptors with the single-residue 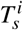 scores.

**Figure 6.**
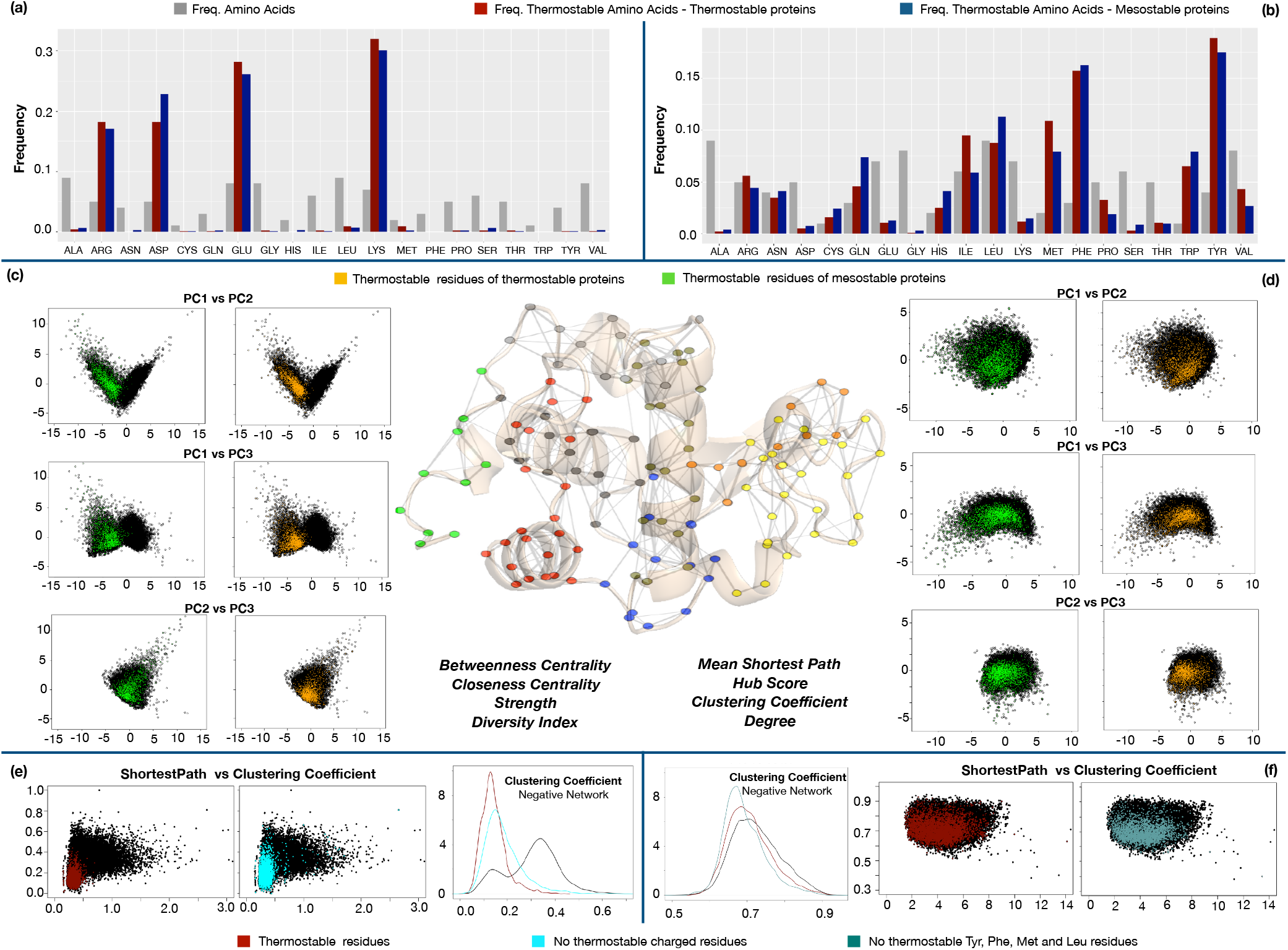
(color online) **a-b)** Frequencies of thermostable amino acids for the thermostable and mesostable proteins are shown in red and blue respectively. Frequencies of all the amino acids are shown in gray. **c–d)** Projection along the first three principal components of all residues. Thermostable residues for mesostable and thermostable proteins are indicated in green and in orange dots, respectively. **e-f)** All residues are mapped in LC space. In red Arg, Asp, Glu and Lys amino acids are shown as the most frequent thermostable residues of the Coulomb network. In yellow dots, Tyr, The, Leu and Met are shown as the most frequent thermostable amino acids identified by van der Waals based *T_s_* score.

## Discussion

Proteins evolved to be functional in very distinct thermal conditions. The mechanisms proteins have developed to face thermal noise have been studied for a long time, given the influence that the comprehension of these mechanisms could exert on both the academic and theoretical industrial field. However, the complete understanding of the reasons that rule the fold stability is a challenging and unsolved problem in which a number of factors have to be taken into account.

Comparative studies of homologous pairs have previously reported a change of content in the amino acids of thermostable proteins^54^ and Arg, Glu and Tyr are more frequently at the surface of thermophilic proteins with Tyr being involved in the formation of stabilizing aromatic clusters^22,27,46^. Undoubtedly, these studies have contributed to unveil mechanisms of thermal resistance typical of specific protein families, yet they do not provide a unifying, global theory describing the rules of adaptation at extreme conditions. In fact, distinct chemical physical characteristics or different combinations of attributes contribute differently to the stabilization of a protein family or fold, making not trivial to use homologous-based findings to infer the thermal stability of a given protein.

At present, just two methods have been developed to predict the melting temperature of a given protein without need of comparison with homologs and relying on a dataset of known *T_m_*. Ku *et al*.^19^ proposed a sequence-based methods of statistical inference that relates the number of dipeptides within the amino acid sequence to the protein *T_m_* allowing one to separate high thermostable from low thermostable proteins. More recently, Pucci et al.^15^ developed a method, based on the thermodynamic statistical potentials, that is able to predict the melting temperature of a given protein using as inputs the three-dimensional structure and the additional information of the optimal temperature (*T_env_*) the protein host organism lives in.

Our work aims to represent a step toward the understanding of the thermal properties of a protein given its 3D structure. While the axiom thermophilic organisms have thermostable proteins is certainly correct, some mesophilic proteins may as well be thermostable^31^. Knowledge on the organism optimal growth temperature, *T_env_*, used to classify mesophiles and thermophiles, may be misleading with high value of correlation due to the fact that *T_env_* is always a lower-bound for *T_m_*.

The basic idea behind our method relies on the assumption that thermostable proteins undergo an optimization process during evolution that leads to specific structural arrangement of their energy interactions. Our analysis is based on a residue interaction network (RIN) in which the three dimensional structure of a protein is schematized as a graph with the residues acting as nodes and the molecular interactions as links. The graph representation is frequently adopted to study complex biological systems involving multiple interacting agent^55–59^.

In our definition of network, links are weighted according to the sum of two nonbonded energetics terms: electrostatic and Lennard-Jones potential. The analysis of the distribution of energies (links) highlighted the correlation between the thermal stability of protein sets (grouped according to their *T_m_*) and the probability of finding high intramolecular interactions, with a highest correlation of 0.90 considering eight groups of proteins (Figure 1).

Unfortunately, neither it is possible to further divide the dataset in more groups due to the dataset dimension, nor we could not consider the energy distribution for the single protein because the small number of links makes the statistics noisy, especially in strong energy regions. Moreover, moving to higher orders of organization, e.g. considering the individual residual energies (Strength parameter), further reduces the data. For this reason, the next-up analysis were performed with a two-groups division of the dataset.

Interestingly, we found that not only strong negative energies determine the thermal stability of a protein, but also strong positive interactions play a role. Such finding confirms the complex nature of the protein interaction network and in fact the stabilizing role of repulsive energies can be explained in cases where repulsion between a couple of residues results in a better spatial rearrangement of protein regions. In order to investigate the complex arrangement of the interactions in the 3D protein structure, we performed an analysis based on graph theory approach aimed at the comprehension of the favorable and unfavourable energies disposition. We determined the stabilizing contribution of each amino acid, defining the Strength of a residue (Eq. 6) as the sum of the energies of all the interactions (favorable and unfavorable) the residue establishes with all other residues. Indeed, this parameter gives an estimate of the residue significance in the overall protein architecture and can be used both as a local property of each individual amino acid and as a global average network feature of the entire protein. Moving to the higher level of organization we investigated the biological role of the secondary structure interactions in thermal stability. The interactions between residues belonging to alpha helixes and loops concentrate more energy in thermostable proteins than mesostable ones. Those results suggest that the thermal stability of a given protein is deeply linked both to the intensity of interactions and to their spatial disposition, and that both are fine-tuned during the evolutionary process. In order to assess the thermal stability, we investigated the network energy organization and compared it against an ensemble of randomized networks. The ensemble comparison has two main purposes: The first consists in overcoming the limitation of the need of pairs of homologous proteins for direct comparison. The second purpose, raised from the observation that thermostable proteins are enriched of high connected nodes (hubs) and have more organized networks of interactions respect mesostable proteins^60–62^, relies in the need introducing a quantitative measure of the evolutionary optimization process thermostable proteins underwent, i.e. the distance between real protein interaction network and a randomized one, in which we disrupt the optimization of energy achieved by thermostable proteins during evolution. As described in the method section, the energies of a network are always obtained from a distribution of mesostable protein interactions. In this way, the more the original network diverts from the ensemble, the higher the probability that the protein belongs to the thermostable class. Moreover, the comparison allows us to assess in a quantitative way the effect of the energetic topology of the protein. Using this protocol to build up the Ts parameter-free descriptor and performing a cluster analysis, we are able to discriminate between mesostable and thermostable proteins, with a maximum accuracy of 76% and an maximum Area Under The Curve (AUC) of 78%.

At last, we investigated whether evolution acts on particular residues to optimise protein thermal stability or if stability is given by a cooperative effect with evolution acting on the whole protein. Our analysis identifies two sets of key (thermostable) residues according to the kind of energetic interactions the network is built with (Coulomb or van der Waals). Surprisingly, thermostable residue frequency in thermostable and mesostable proteins is comparable and they represent only a small subset of all residues. In order to better understand the theoretical aspects of thermostability and improve the classification to be used in more applicative fields, we created a new parameter dependent *T_s_* score given by a linear combination of the *T_s_* score of the eighth possible set of residues (see Results). The improved performance of 83% of ROC’s AUC highlighted the promising features of the single residue approach.

## Methods

### Datasets

Proteins with known melting temperature (*T_m_*) were obtained from the ProTherm database^63^. We selected all wild-type proteins for which the following thermodynamic data and experimental conditions were reported: *T_m_* ≥ 0 *^o^C*; 6.5 ≤ pH ≤ 7. 5 and no denaturants. Experimentally determined structures were collected from the PDB^34^and filtered according to method (x-ray diffraction), resolution (below 3Å) and percentage of missing residues (5% compared to the Uniprot^64^ sequence). Proteins for which experimentally determined structures were only available in a bound state, i.e. in complex with either a ligand or a ion, were excluded. Proteins were filtered using the CD-HIT software^65^ to remove proteins with chain sequence identity ≥ 40% to each other. The final dataset, hereinafter referred to as the *T_m_* dataset, consisted of 71 proteins. Consistently with previous reported dataset, thermostable proteins (*T_m_* ≥ 70*^o^C*) represent about a third of the overall dataset^46,66,67^. In order to have a dataset as balanced as possible, we also manually collected a second, independent dataset consisting of proteins from hyperthermophilic organisms with optimal growth at T ≥ 90 *^o^C* and pH between 6.5 and 7.5. Experimentally determined structures were collected and filtered according to same criteria described above for the Tm dataset, leading to a total of 13 protein structures. This second dataset is referred to as the *T_hyper_* dataset. The union of the two dataset, referred as the *T_whole_* dataset, accounts of 84 proteins (see Supplementary Information).

CATH class and architecture for protein domains were checked: the most representative class for thermostable domains is Alpha Beta (84% respectively) with only a few Mainly Beta domains (11%) and just 1 Mainly Alpha domain (0.02%). On the other hand, Mainly Alpha and Mainly Beta constitutes the 21% and 26% of the mesostable domains, with Alpha Beta sets to 53%. 15 different folds are available for mesostable proteins, with the most representative one being 3-Layer(aba) Sandwich (19%) while the thermostable domains count 9 different folds, with 31% of 2-Layer Sandwich and 3-Layer(aba) Sandwich.

Protein structures were minimized using the standard NAMD^68^ algorithm and the CHARMM force field^69^ in vacuum. A 1 fs time step was used and structures were allowed to thermalize for 10000 time steps.

### Structural analysis

Proteins from both the *T_m_* and *T_hyper_* datasets were analyzed for their secondary structure content and architecture according to the CATH Protein Structure Classification database^70^. Per residue secondary structure assignment was done using the DSSP software^71^. In order to assess how the energy is distributed among protein secondary structure elements we assigned each couple of residues to a class of interaction, based on which secondary structure element they belong to. Possible interaction classes are: helix-helix, helix-strand, helix-loop, strand-strand, strand-loop and loop-loop. To evaluate the fraction of total energy proteins devolve to each class we defined the difference between the fraction of observed energy and a theoretical fraction:

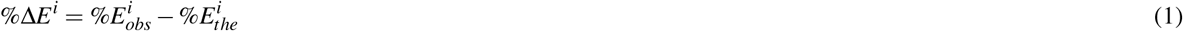

where i represents the pair of secondary structures considered (e.g. helix-helix), 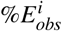 is the ratio between the energy of class i and the total energy 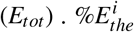 estimates the expected fraction of energy for class i assuming equivalent distribution of energy among classed:

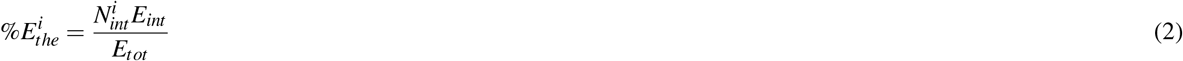

where 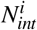 is the number of interactions of class i and *E_int_* is the average energy value. In other words Eq. 2 gives the ratio between the number of interactions of class i and the total number of interactions. Energy distribution densities were calculated using the R density function with default parameters.

### Network representation and analysis

In the present work, protein structures are represented as Residue Interaction Networks (RINs), where each node represents a single amino acid *aa_i_*. The nearest atomic distance between a given pair of residues *aa_i_* and *aa_j_* is defined as *d_ij_*. Two RIN nodes are linked together if *d_ij_* ≤ 12 Å ^68,69^. Furthermore links are weighted by the sum of two energetic terms: Coulomb (C) and Lennard-Jones (LJ) potentials. The C contribution between two atoms, *a_l_* and *a_m_*, is calculated as:

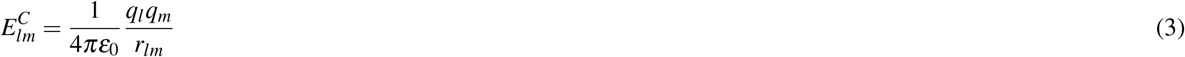

where *q_l_* and *q_m_* are the partial charges for atoms *a_l_* and *a_m_*, as obtained from the CHARMM force-field: *r_lm_* is the distance between the two atoms, and *ε*_0_ is the vacuum permittivity. The Lennard-Jones potential is instead given by:

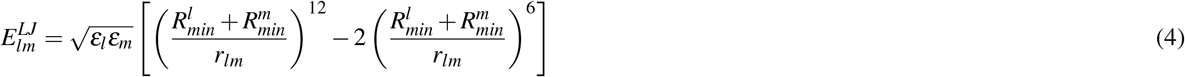

where *ε_l_* and *ε_m_* are the depths of the potential wells of atom l and m respectively, 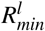 and 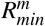 are the distances at which the potentials reach their minima. Therefore, the weight of the link connecting residues *aa_i_* and *aa_j_* is calculated by summing the contribution of the single atom pairs as reported in equation 5.

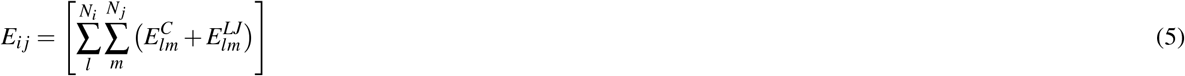

where *N_i_* and *N_j_* are the number of atoms of the i-esime and j-esime residue respectively.

Network analysis has been performed using the igraph package^72^ implemented in R^73^. For each RIN, the Strength local parameter^74^ is defined as:

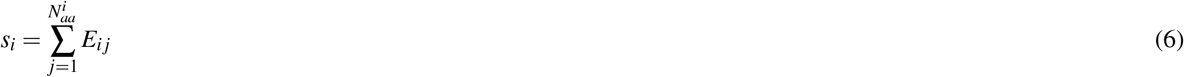

where the Strength *s_i_* of the i-esime residue is calculated as the sum of all energetic interactions for that residue 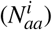. The 3D images of the protein networks were generated using Pymol^75^.

### Network randomization

In order to distinguish mesostable from thermostable proteins, we compare the Strength calculated in the real network against the same parameter obtained from an ensemble of random RINs. More specifically, the Strength of each real network is compared against a distribution of mean Strength values from 500 randomized networks obtained from the real one using the procedure described below.

Given a RIN link characterize by an energy weight *E_ij_* and an interaction distance *d_ij_*, we replace the energy value with a new one 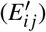, randomly extracted from the energy distribution observed in mesostable proteins from the *T_m_* dataset and in the same distance interval. Given the global range of interaction distances 0-12 *Å*, twenty-four consecutive, non-overlapping distance intervals are obtained by dividing the entire range into a grid of bins using a bin width of 0.5 *Å*. A *T_s_* score, defined as:

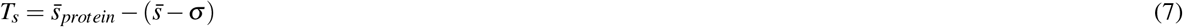

is calculated to estimate how much the original RIN mean Strength value deviates from the expected mean value of rRIN distribution. 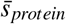 is the average of the Strength parameter for the RIN; 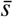 and *σ* are the mean and standard deviation of the average values of the rRIN distribution. At the level of single residue, we define a 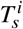 score, similarly to the one in Eq. 7, as

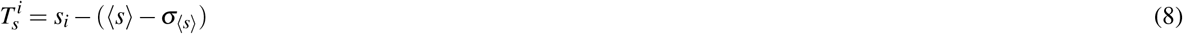

where *s_i_* represents the Strength of residue *i*, 〈*s*〉 is the average Strength of residue *i* over the 500 randomized networks and *σ*_〈*s*〉_ is the standard deviation.

### Performance evaluation

We evaluated the performance of the *T_s_* score in discriminating between termostable and mesostable proteins by a seven cross validation. The 49 mesostable proteins of the *T_m_* dataset were divided in seven groups, guaranteeing that number of residues and *T_m_* values were as broad distributed as possible. For each group of mesostable proteins:

1. twenty-four density distribution 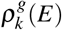 are built, where *g* indicates the groups out of the seven created and E stands for the total energy defined in Eq. 5.
2. The remaining 42 mesostable proteins and the 22 thermostable ones together with the *T_hyper_* dataset proteins, are randomized according to previous described procedure sampling the weights from the 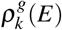.
3. All randomized proteins are classified as mesostable or thermostable proteins according to the obtained *T_s_* score.

This procedure ensures that the classification of the mesostable proteins is not biased by their own presence in the energy density distributions used in the randomization process. We used the R package pROC^76^ to plot the ROC curve and calculate the AUC values.

### Clustering analysis

We clustered the *T_s_* descriptors using the Euclidean distance and the Ward method as linkage function^77^ via the “hclust” function of the “Stats” package of R^73^. To better compare the different *T_s_* score between them we normalize the data dividing each *T_s_* score for the maximum of the absolute values. Finally, using the R package “clValid”^78^, we performed an internal validation for the hierarchical cluster considering both the Connectivity, Dunn and Silhouette parameters.

### Principal Component Analysis

PCA was performed over eight graph-based descriptors using “princomp” function of R software and the correlation matrix was used for the analysis^79^. Each descriptor has been computed using a specific function available in the R i-graph package. The involved descriptors and corresponding functions are: Betweenness Centrality *(betweenness* function)^80^; Closeness Centrality (*closeness* function)^80^; Strength, (*strength* function)^74^; Diversity Index, *(diversity* function)^81^; Mean Shortest Path, (*distances* function, with “dijkstra” algorithm)^82^; Hub Score, (*hubscore* function)^83^; Clustering Coefficient, (*transitivity* function, with “barrat” algorithm)^74^; Degree, (*degree* function)^82^.

## Acknowledgements

The authors dedicate this article to the memory of Professor Anna Tramontano, whose striking ideas lied the basis of the present work.

The research leading to these results has been supported by Epigenomics flagship project EPIGEN. G.G.T. is funded by European Research Council (*RIBOMYLOME*_309545), Spanish Ministry of Economy and Competitiveness (BFU2014 – 55054 – *P* and *BFU*2017 – 86970 – P) and “Fundació La Marató de TV3” (PI043296).

## Author contributions statement

E.M. and R.L. conceived the study. G.G.T. contributed with additional ideas. M.M. and E.M. performed the calculations, M.M., P.P.O., L.D.R., F.A., P.C. and G.G.T. analysed the data. P.P.O. and F.A. collected the dataset. E.M., M.M., R.L. and G.G.T. wrote the manuscript. All authors reviewed the manuscript.

